# Selection of internal references for transcriptomics and RT-qPCR assays in Neurofibromatosis type 1 (NF1) related Schwann cell lines

**DOI:** 10.1101/2020.10.22.350017

**Authors:** Yi-Hui Gu, Xi-Wei Cui, Jie-Yi Ren, Man-Mei Long, Wei Wang, Cheng-Jiang Wei, Rehanguli Aimaier, Yue-Hua Li, Man-Hon Chung, Bin Gu, Qing-Feng Li, Zhi-Chao Wang

## Abstract

Transcriptomics has been widely applied in uncovering disease mechanisms and screening potential biomarkers. Internal reference selection determines the accuracy and reproducibility of data analyses. The aim of this study was to identify the most qualified reference genes for the relative quantitation analysis of transcriptomics and real-time quantitative reverse-transcription PCR in fourteen NF1 related cell lines, including non-tumor, benign and malignant Schwann cell lines. The expression characteristics of eleven candidate reference genes (RPS18, ACTB, B2M, GAPDH, PPIA, HPRT1, TBP, UBC, RPLP0, TFRC and RPL32) were screened and analyzed by four software programs: geNorm, NormFinder, BestKeeper and RefFinder. Results showed that GAPDH, the most used internal reference gene, was significantly unstable. The combinational use of two reference genes (PPIA and TBP) was optimal in malignant Schwann cell lines and the use of single reference genes (PPIA or PRLP0) alone or in combination was optimal in benign Schwann cell lines. Our recommended internal reference genes may improve the accuracy and reproducibility of the results of transcriptomics and real-time quantitative reverse-transcription PCR in further gene expression analyses of NF1 related tumors.

## Introduction

Neurofibromatosis type 1 (NF1) is an autosomal dominantly inherited tumor predisposition syndrome that affects multiple organ systems and has a wide range of variable clinical manifestations, such as pigmentary lesions, skeletal abnormalities[1–3]. The major defining features in NF1 patients are peripheral nerve sheath tumors, including plexiform neurofibromas (pNF) and malignant peripheral nerve sheath tumors (MPNST)[4]. Therefore, the majority of NF1 related tumor researches could be categorized into benign tumor studies or malignant tumor studies.

Currently, Omics studies play an increasingly important role in uncovering disease mechanisms and screening potential therapeutic biomarkers. Among different omics studies, transcriptomics is the bridge between genomics and proteomics. Key methods, including microarray and RNA-seq, could provide deep and precise measurements of levels of transcripts and different isoforms. Several studies are conducted to figure out the molecular mechanisms of NF1 and to advance the development of targeted drugs[5–7]. These high-throughput results need further validation by real-time quantitative reverse-transcription PCR (RT-qPCR), which is considered as the gold standard for gene expression studies. However, the accuracy and reproducibility of these results may be affected by several factors at multiple stages. In data analysis, the stability of reference genes is critical for appropriate standardization and obtaining accurate gene expression data. The Minimum Information for Publication of Quantitative Real-Time

PCR experiments (MIQE) guidelines highlight the importance of experimental validation of reference genes for particular tissues or cell types[8–11]. However, to date, there is no previous research on the validation of suitable reference genes for the relative quantification analysis of target gene expression in NF1 related cell lines.

In the present study, eleven candidate reference genes, which are most frequently used for relative quantification analysis in other neoplastic diseases, were selected, including 18S Ribosomal RNA (RPS18), β-actin (ACTB), β-2-microglobulin (B2M), Glyceraldehyde-3-phosphate dehydrogenase (GAPDH), Peptidylprolyl isomerase A (PPIA), Hypoxanthine phosphoribosyltransferase 1 (HPRT1), TATA binding protein (TBP), Ubiquitin C (UBC), Ribosomal protein large P0 (RPLP0), Transferrin receptor (P90, CD71) (TFRC), and Ribosomal protein 32 (RPL32)[12–16].

We investigated the stability of these eleven reference genes in fourteen different NF1 related cell lines, including two non-tumor *NF1^+/-^* Schwann cell lines, five benign pNF cell lines and seven malignant MPNST cell lines. Based on study design, we analyzed the data obtained as two separate groups: (1) benign NF1 tumor study including Schwann cell (SC) + pNF cell lines and (2) malignant NF1 tumor study including SC + pNF + MPNST cell lines, aiming to identify the most qualified reference genes for gene expression analysis in benign and malignant NF1 tumor study respectively.

## Materials and Methods

### Neurofibromatosis type 1 (NF1) related cell lines

Five MPNST cell lines (ST-8814, STS26T, S462, S462TY, T265) were generous gifts from Dr.Vincent W Keng (Department of Applied Biology and Chemical Technology, The Hong Kong Polytechnic University, Kowloon, Hong Kong) and Jilong Yang (International Medical School, Tianjin Medical University, Tianjin). Two pNF cell lines (A68 and WZJ) are immortalized as described previously[17]. Ethics Committee of Shanghai Ninth People’s Hospital, Shanghai Jiao Tong University School of Medicine approved this study. Informed consent was achieved from patients under institutional reviewer board protocols. hTERT NF1 ipNF05.5 (ATCC^®^ CRL-3388^TM^), hTERT NF1 ipNF95.6 (ATCC^®^ CRL3389™), hTERT NF1 ipNF95.11b (ATCC^®^ CRL3390™), hTERT NF1 ipnNF95.11c (ATCC^®^ CRL3391™), hTERT NF1 ipn02.3 2λ (ATCC^®^ CRL3392 ^TM^), sNF96.2 (ATCC^®^ CRL2884 ^TM^) and sNF02.2 (ATCC^®^ CRL2885 ^TM^) were obtained from the American Type Culture Collection (ATCC). All the cell lines used in this study were cultivated in Dulbecco’s Modified Eagle Medium (DMEM) supplemented with 10% fetal bovine serum (FBS) and 1% penicillin/streptomycin at 37°C in a humidified incubator at 5% CO2 and were confirmed negative for mycoplasma prior to use. Verification of cell lines was performed by Short Tandem Repeat (STR) DNA profiling (Applied Biological Materials Inc, Canada).

### Total RNA extraction and cDNA synthesis

Total RNA was extracted from all NF1 related cell lines using AxyPrep Multisource RNA Miniprep Kit (Axygen, USA) according to the manufacturer’s instructions. The quality and quantity of RNA were measured using a NanoDrop 2000 Spectrophotometer (Thermo Scientific, Waltham, MA, USA) through OD260/280 and OD260/230 ratios. 500 ng of total RNA was used for the cDNA synthesis reaction using PrimeScript™ RT Master Mix kit (TaKaRa RR036A) according to the manufacturer’s instructions. After the reaction, the cDNA was diluted to 20 ng/ml. All RT-qPCR experiments were performed with the same batch of cDNA.

### RT-qPCR

Reference gene primers were designed and synthesized by Sangon Biotech (Shanghai, China). The efficiency, dynamic range and specificity of all primers were tested by Sangon Biotech (Shanghai, China). RT-qPCR was performed using TB Green^®^ Premix Ex Taq™ Kit (TaKaRa RR420A) on a Applied Biosystems 7500 Real-Time PCR System, as described previously[18]. Each sample was measured in three technical replicates. The relative quantification of RT-qPCR data were calculated using the 2^-ΔΔC^_T_ method, as described previously[8].

### Statistical analysis

The stability of the eleven candidate reference genes was examined by four frequently used software programs, geNorm[19], NormFinder[20] BestKeeper[21] and RefFinder[22], as described previously[23–26].

## Results

### The expression characteristics of eleven internal reference gene candidates

The threshold cycle (Ct) value was used to assess the expression characteristics of the internal reference gene candidates. Higher Ct values indicate lower expression levels. The distribution of the C_t_ values of eleven reference genes in fourteen samples, including two *NF1^+/-^* Schwann cell lines, five plexiform neurofibroma cell lines and seven MPNST cell lines was displayed in Fig 1. The C_t_ values represented in this research from all samples ranged from 18.09 to 43.65. It is worth noting that UBC showed significantly higher C_t_ values in MPNST samples compared to other samples, suggesting lower mRNA expression level.

**Fig 1.**
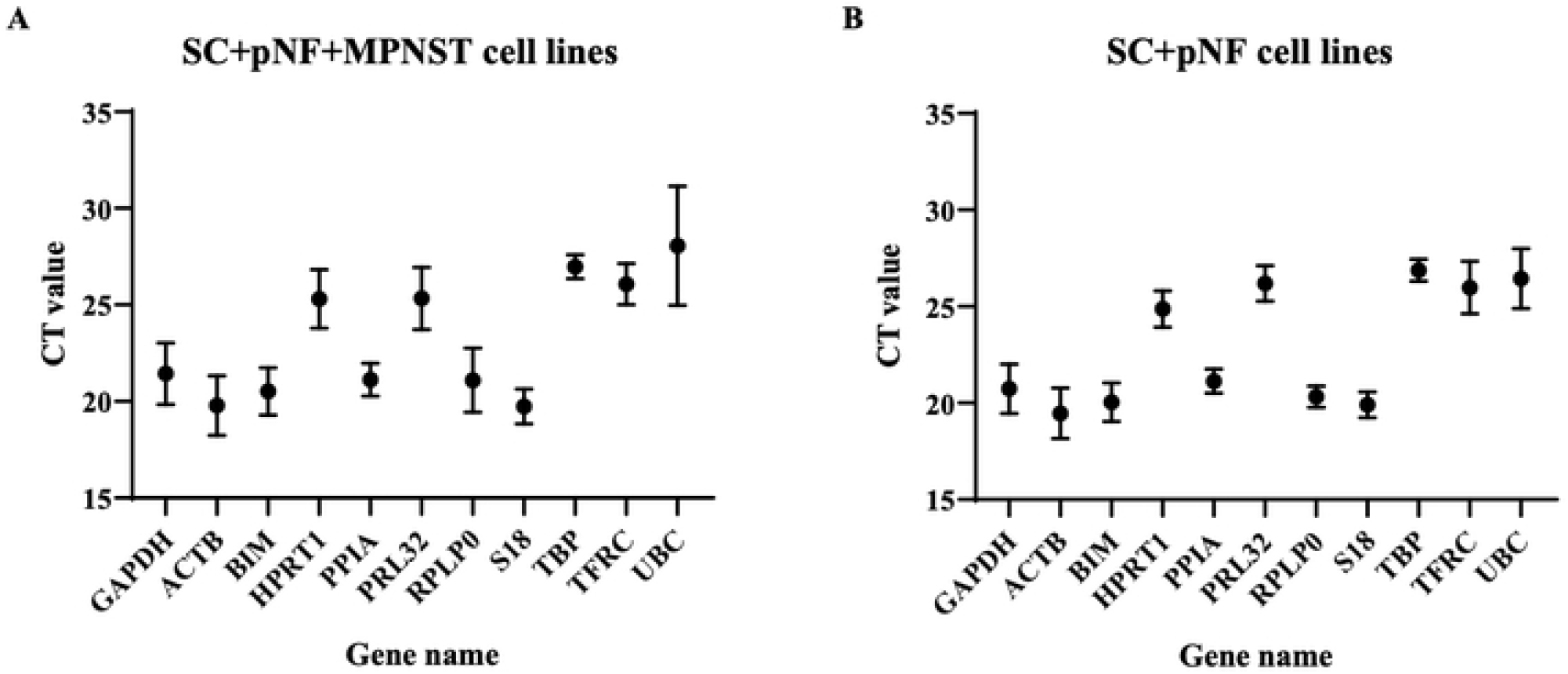
C_t_ values of the candidate internal reference genes. Dots represent the mean C_t_ value; bars represent the mean ± standard deviation. (A) C_t_ values of each candidate internal reference gene in *NF1*^+/-^ SC, *NF1*^-/-^ pNF and MPNST cell lines (*n*=14). (B) C_t_ values of each candidate internal reference gene in SC and pNF cell lines *(n* = 7). SC, Schwann cell; pNF, plexiform neurofibroma; MPNST, malignant peripheral nerve sheath tumor.

### Genorm analysis

In order to identify the best reference genes for benign and malignant NF1 tumor study, we applied four statistical approaches including geNorm, NormFinder, BestKeeper and ΔC_t_ method. The *M* value of eleven reference genes were calculated and ranked by genorm. The genorm system identifies those genes with lowest *M* values as the most stably expressed genes and an *M* value under 0.5 represents relatively stable gene expression. GeNorm analysis showed that PPIA and TBP, sharing an *M* value of 0.549, were the most stable reference genes in SC + pNF + MPNST cell lines (Fig 2). However, when only considering SC + pNF cell lines, the most stable genes were PPIA and PRLP, with an *M* value of 0.225, followed by TBP and S18 (*M* value, 0.295 and 0.454, respectively). The optimal number of reference genes to obtain a stable normalization index was demonstrated by V-values. When SC + pNF + MPNST cell lines were considered, the V8/9 value was 0.127, indicating that eight reference genes combination is optimal for RT-qPCR normalization. In SC + pNF cell lines, the V2/3 value was 0.104, suggesting that merely two reference genes were necessary. And the V8/9 value was lower than 0.110 in this group, of which the V values kept stable in different subsets.

**Fig 2.**
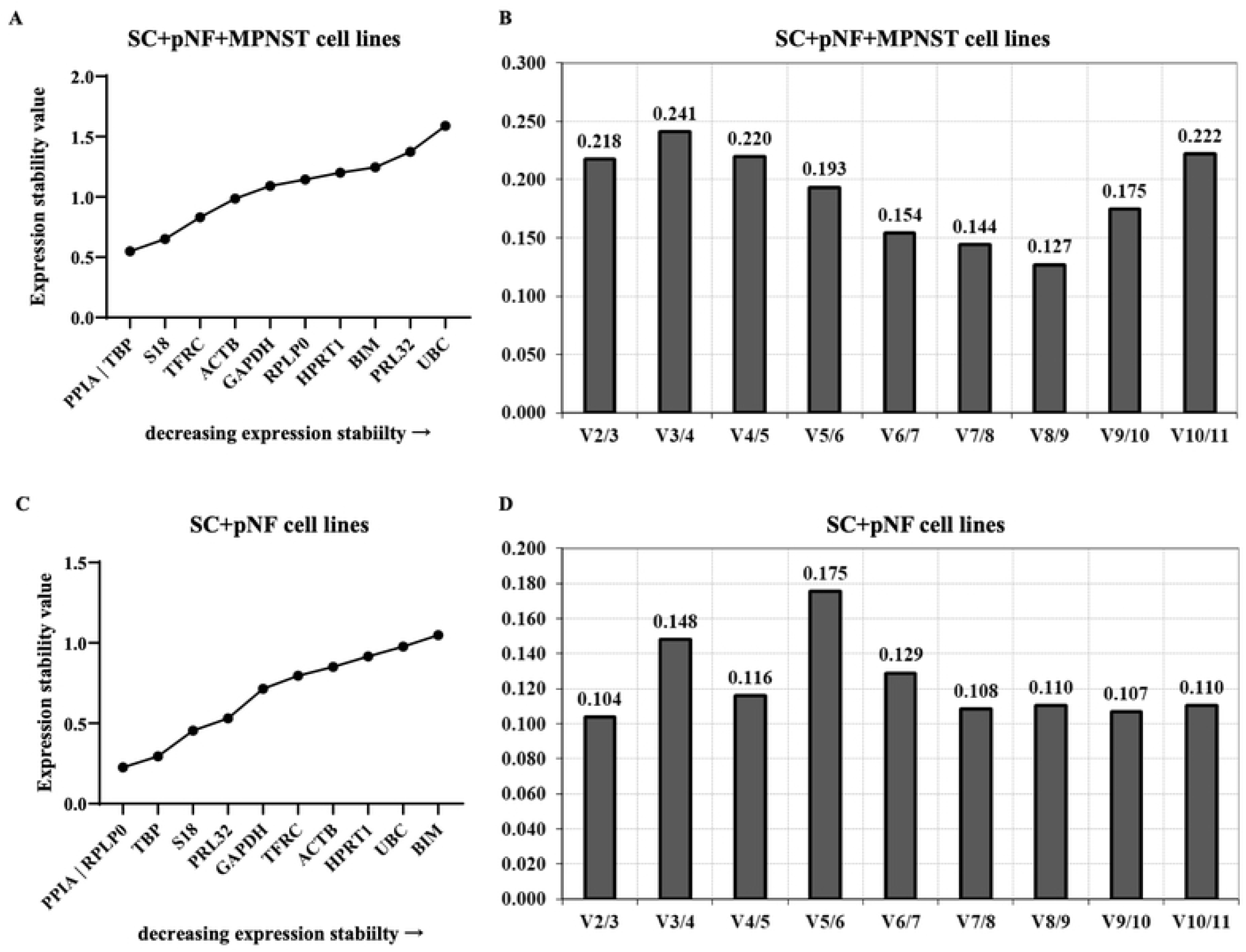
Analysis results of geNorm program. The expression stability of eleven candidate genes in SC + pNF + MPNST subset (A) and SC + pNF subset (C) was calculated and ranked separately. The *x-axis* represents various candidate reference genes, and the *y-axis* represents stability value (*M* value). Lower *M* value suggests higher expression stability. B and D show the optimal number of reference genes in different subsets. The *x-axis* represents the number of genes selected for comprehensive analysis *V* (*n/n*+1), and the *y-axis* means the pairwise variation value (*V* value). When *V* value is under 0.15, the corresponding combination is esteemed stable and *n* is the best number of internal reference genes. SC, Schwann cell; pNF, plexiform neurofibroma; MPNST, malignant peripheral nerve sheath tumor.

### NormFinder analysis

Eleven reference genes underwent normalization factor calculation and were ranked by NormFinder according to their minimal combined SC, pNF and MPNST gene expression variation. According to the result, TBP was the optimal reference gene (stability value, 0.394), followed by PPIA, ACTB and GAPDH (stability values, 0.403, 0.427 and 0559 respectively) (Fig 3). The best combination of two genes are GAPDH and S18 with a combination stability value of 0.150. For SC + pNF cell lines, PPIA, RPLP and TBP were the most stably expressed genes (stability values, 0.142, 0.185 and 0.186 respectively), while the rest ones were extremely unstable amongst those cell lines.

**Fig 3.**
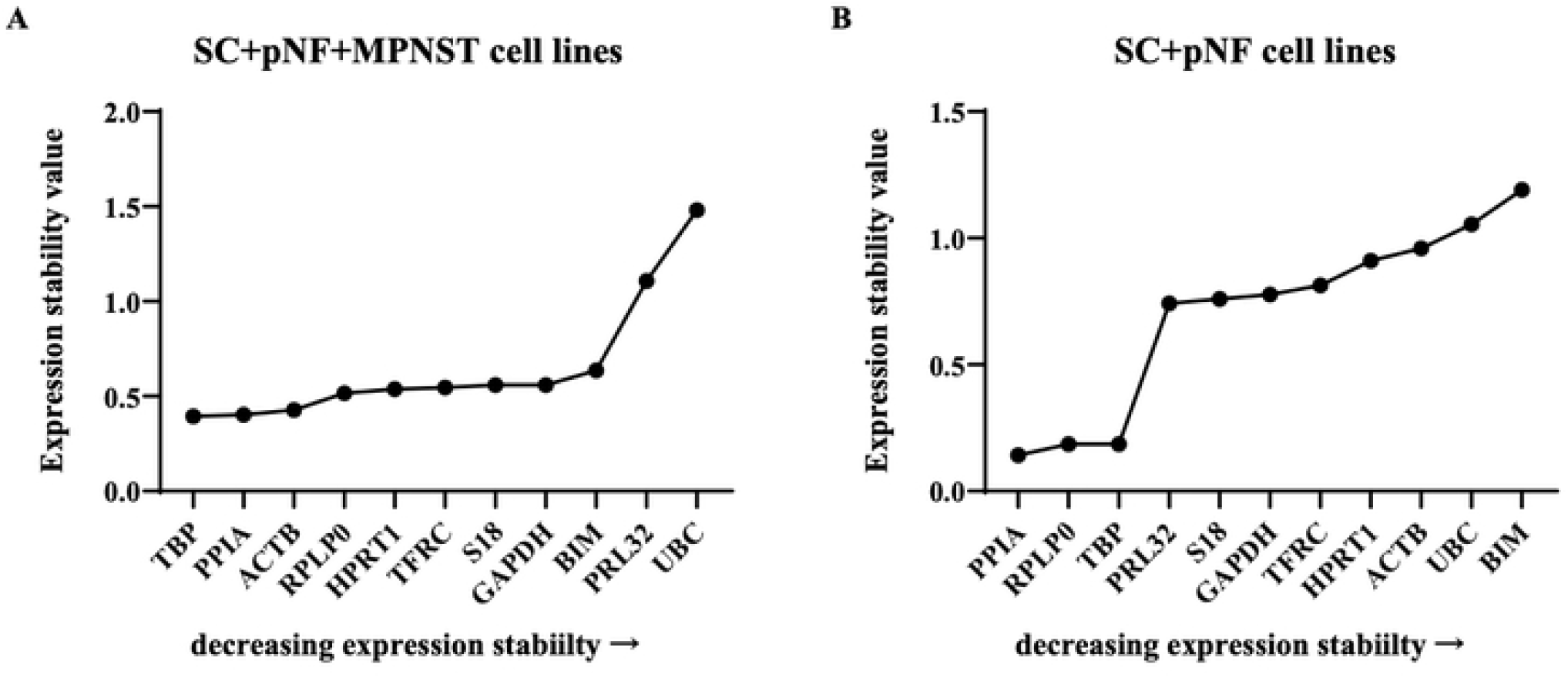
Analysis results of NormFinder program. The *x-axis* represents various candidate reference genes, and the *y-axis* represents stability value. (A) The stability value of each candidate internal reference gene in SC + pNF + MPNST subset (*n*=14). (B) The stability value of each candidate internal reference gene in SC + pNF subset (*n*=7). SC, Schwann cell; pNF, plexiform neurofibroma; MPNST, malignant peripheral nerve sheath tumor.

### Best keeper analysis

In best keeper algorithm, the C_t_ values, SD (standard deviation) and CV (coefficient of variance) of each gene were calculated and analyzed to identify stable reference gene candidates. Under general conditions, genes with a SD greater than 1.0 are determined to be unstable. Among SC + pNF + MPNST cell lines, TBP turned out to be the best choice with a standard deviation [+/− CP] (which is the stability value in best keeper algorithm) of 0.48 (Fig 4). The stability value of TBP kept stable (0.44) when it came to SC + pNF cell lines. However, RPLP and PPIA were esteemed as more stable genes, sharing a stability value of 0.42.

**Fig 4.**
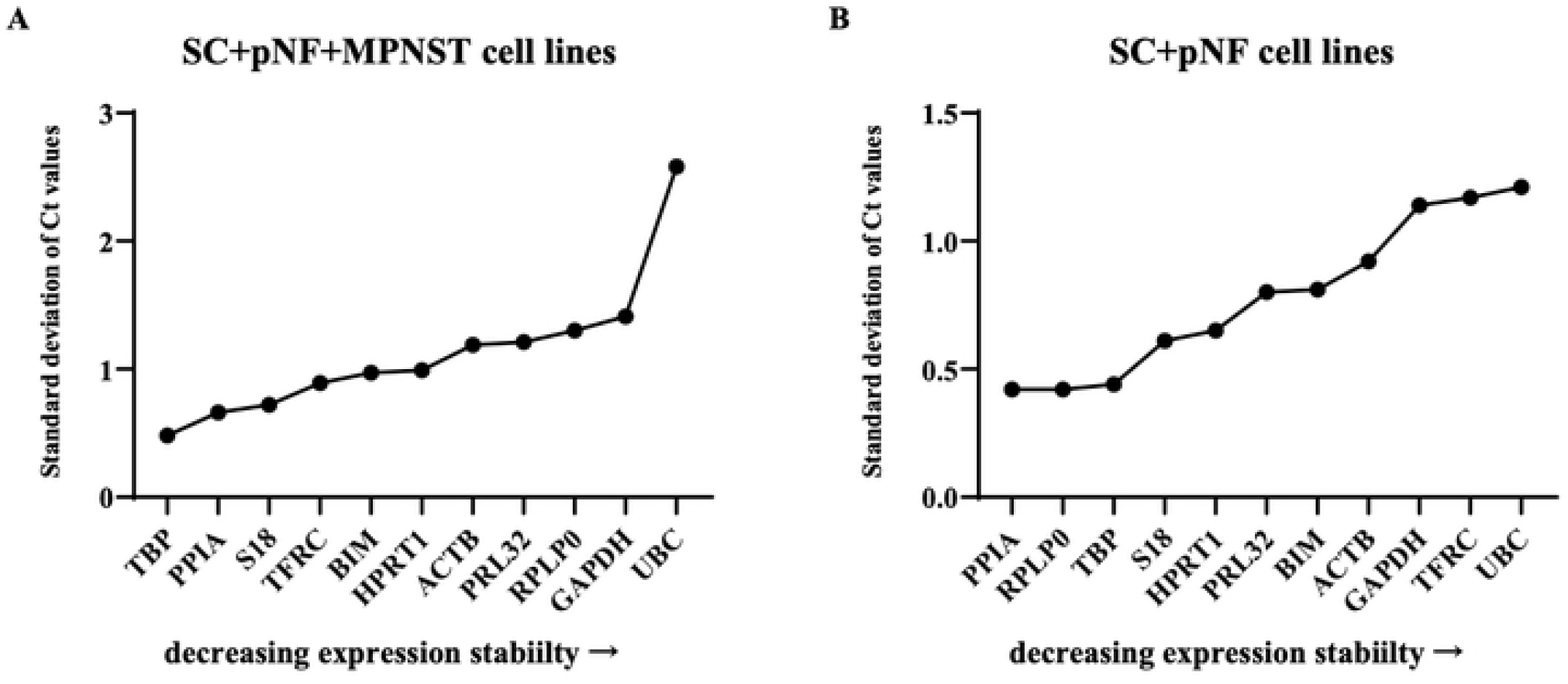
Analysis results of BestKeeper program. The *x-axis* represents various candidate reference genes, and the *y-axis* represents stability value. (A) The stability value of each candidate internal reference gene in SC + pNF + MPNST subset (*n*=14). (B) The stability value of each candidate internal reference gene in SC + pNF subset (*n*=7). SC, Schwann cell; pNF, plexiform neurofibroma; MPNST, malignant peripheral nerve sheath tumor.

### ΔC_t_ analysis

In ΔC_t_ method, the differential expression of “gene pairs” were analyzed to determine the optimal reference genes. According to the results, PPIA (1.24) and TBP (1.26) were the most stably expressed reference genes in SC + pNF + MPNST cell lines (Fig 5). While RPLP (0.78) was far more stable in SC + pNF cell lines.

**Fig 5.**
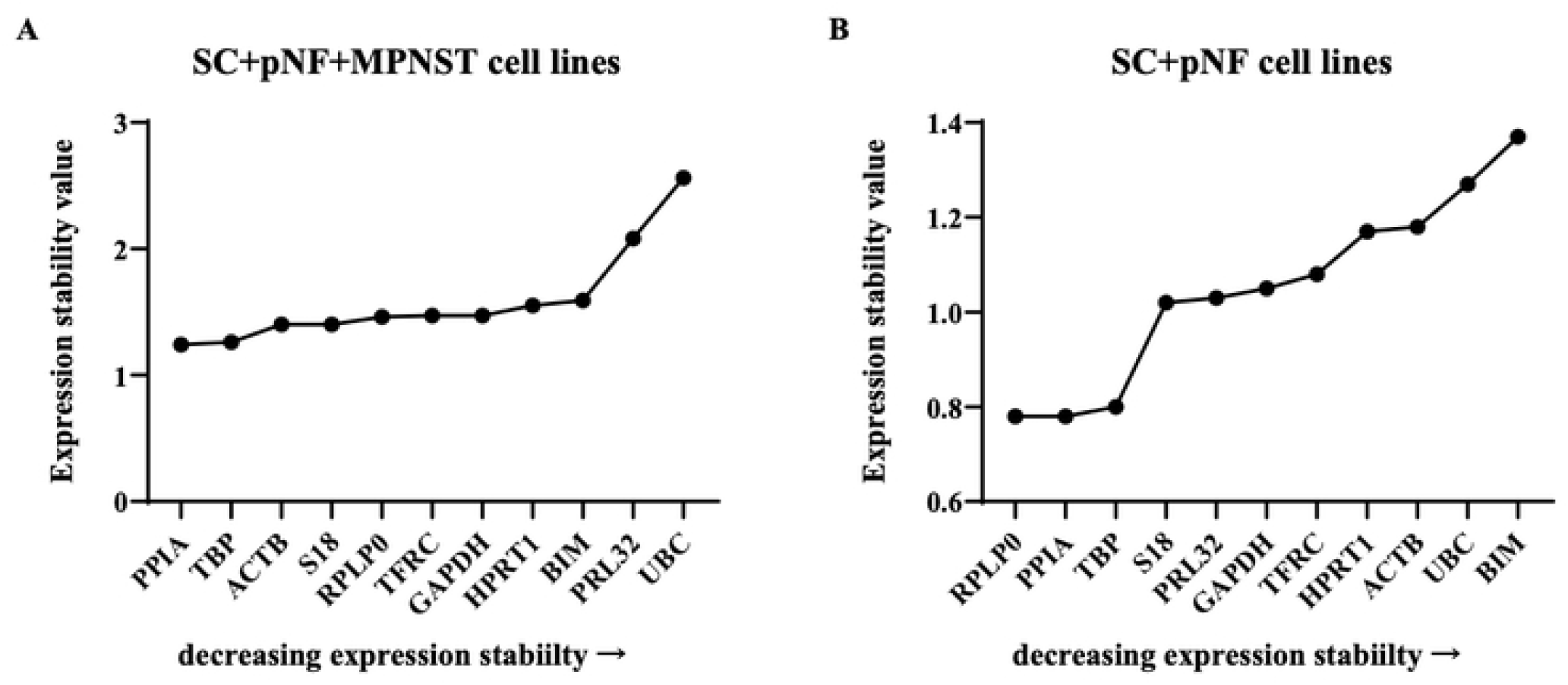
Analysis results of ΔC_t_ algorithm. The *x-axis* represents various candidate reference genes, and the *y-axis* represents stability value. (A) The stability value of each candidate internal reference gene in SC + pNF + MPNST subset (*n*=14). (B) The stability value of each candidate internal reference gene in SC + pNF subset (*n*=7). SC, Schwann cell; pNF, plexiform neurofibroma; MPNST, malignant peripheral nerve sheath tumor.

### Comprehensive ranking order

Using RefFinder, we integrated four analysis approaches mentioned above to comprehensively evaluate the expression stability of these candidate genes. PPIA turned out to be the optimal reference gene in SC + pNF + MPNST cell lines with a ranking value of 1.19, followed by TBP and S18 (ranking value, 1.41 and 3.66, respectively) (Fig 6). And PRLP (1.19) was the most stably expressed in SC + pNF cell lines.

**Fig 6.**
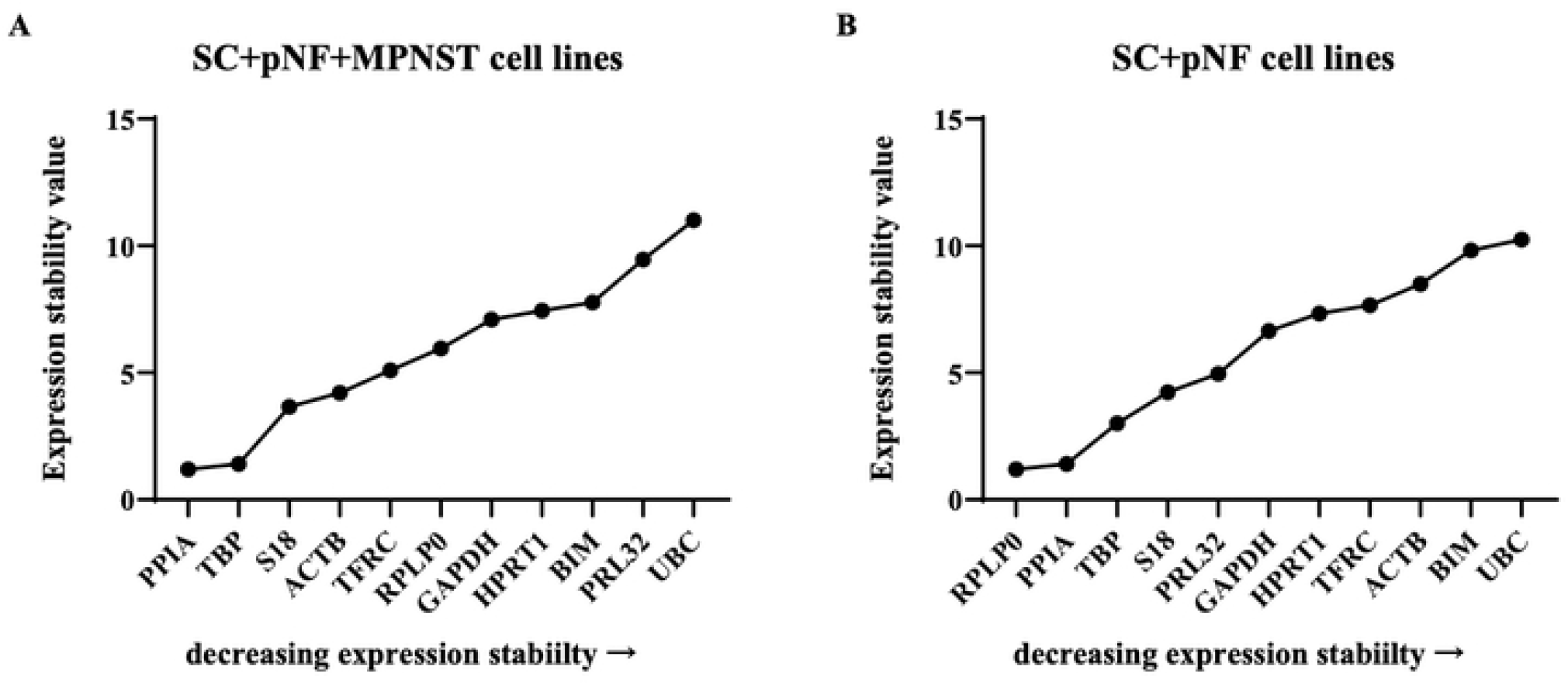
Analysis results of ReFinder program. The *x-axis* represents various candidate reference genes, and the *y-axis* represents stability value. (A) The stability value of each candidate internal reference gene in SC + pNF + MPNST subset (*n*=14). (B) The stability value of each candidate internal reference gene in SC + pNF subset (*n*=7). SC, Schwann cell; pNF, plexiform neurofibroma; MPNST, malignant peripheral nerve sheath tumor.

## Discussion

High-throughput transcriptome sequencing identifies the key pathogenic genes that play crucial roles in the pathogenesis, development and malignant transformation of NF1 related neurofibroma. RT-qPCR has been considered as golden standard for gene expression analysis because of its accuracy and sensitivity[27]. It is a robust and specific method for the validation of the identity of these pathogenic genes that aberrantly expressed in neurofibroma.

The ease to generate RT-qPCR data is in sharp contrast with the challenges to guarantee that the results obtained are reliable. In the absence of proper reference genes, data obtained are possibly inaccurate and unreproducible, as it has been shown that the use of a single reference gene without validation results in a significant bias (ranging from 3-fold in 25% of the results up to 6-fold in 10% of the results)[19].

Prior to this study, no validated reference gene has been identified for NF1 related cell lines. In previous studies, GAPDH and ACTB have been most frequently used in gene expression analysis of NF1 without solid validation[28–30]. However, the expression characteristics of reference genes vary remarkably under different experimental conditions and with different samples. The use of GAPDH and ACTB as an internal control has been proven to be unsuitable in other samples, such as lymphoblastoid cell lines and human mesenchymal stem cells[31–33]. Therefore, it is crucial to confirm reliable and qualified reference genes for particular tissues or cell types and specific experimental designs.

In our study, in order to find the internal reference genes stably expressed in different NF1 samples, including normal peripheral nerve, benign and malignant tumor tissues, we investigated fourteen different NF1 related cell lines, derived respectively from the tissues mentioned above, which include two non-tumor *NF1^+/−^* Schwann cell lines, five pNF cell lines and seven MPNST cell lines. The expression stability of eleven frequently used reference genes, including RPS18, ACTB, B2M, GAPDH, PPIA, HPRT1, TBP, UBC, RPLP0, TFRC, and RPL32 were investigated and analyzed separately in these cell lines.

Four different mathematical and statistical models were utilized to analyze the data obtained, including geNorm, NormFinder, BestKeeper and ΔC_t_. Each model uses different algorithms to estimate both the intra- and the intergroup expression variations and rank candidate genes based on the instability score. GeNorm and NormFinder use the stability (actually instability) value, ΔC_t_ uses the average of S.D. and BestKeeper uses the S.D. of the crossing points[19–21]. In addition, A web-based tool RefFinder was finally used to evaluate the stability of candidate reference genes and identify the most stable gene by calculating the geometric mean of ranking values obtained from the above-mentioned four methods[22].

Due to the fact that the software applied are based on different mathematical models, the ranking orders of reference gene stabilities varied slightly between these tools. However, the top two positions of reference genes in SC + pNF group and SC + pNF + MPNST group determined by geNorm were identical to those determined by NormFinder, BestKeeper and ΔC_t_. Unanimously, according to the results of RefFinder, the top two reference genes in two groups are also the same and they share similar ranking values which are significantly lower than others. Therefore, conclusion can be drawn that PPIA and PRLP0 are the best reference genes for normalization in benign NF1 tumor study, while PPIA and TBP are as well in malignant NF1 tumor study.

PPIA is a gene encoding for a cyclosporin binding-protein, TBP is a gene encoding for a TATA-binding protein, and PRLP0 encodes a ribosomal protein that is a component of the 60S subunit. In NF1 related tumors, they were stably expressed irrespective of tumor pathology and severity. However, the most frequently used reference gene GAPDH was proved to be an inappropriate option in two groups (*M* values > 0.5). As a multifunctional gene, the use of GAPDH as a reference gene has been also questioned and challenged in other cancers including lung cancer[34], breast cancer[35] and bladder cancer[36]. Accumulated evidences indicate that GAPDH is deregulated in various cancers under certain conditions and potentially participates in tumorigenesis and tumor progression[34–38].

In addition, considering the difference of opinion on the minimal number of reference genes required for RT-qPCR, we explored the necessity of selecting multiple reference genes for data normalization in our samples. In previous studies, some investigators showed that the combination of more than one reference gene improved the accuracy of results[14, 19, 39–41] while other investigators proved that normalization with a single gene is sufficient for most research applications[12, 36, 42, 43]. In present study, according to the analyses of geNorm, combination of eight reference genes is the most precise plan for normalization in SC + pNF + MPNST group. However, considering the stability and precision of using two reference genes in combination is sufficient for data normalization, we suggest using a normalization factor calculated by the geometrical mean of the most stable reference genes (PPIA and TBP) for normalization of target gene expression in SC, pNF and MPNST cells[44]. What’s more, it is worth noting that the use of single reference gene, with a M-value over 0.5, should be avoided in these samples. In SC + pNF group, although the combination of two reference genes significantly improved precision over normalization with PPIA or PRLP0 alone, the use of single reference genes (PPIA or PRLP0), sharing an *M* value of 0.225, is acceptable for data normalization.

## Conclusions

In this study, we systematically explored the suitability of fourteen candidate reference genes for normalization of gene expression in different NF1 related cell lines, including non-tumor *NF1*^+/−^ Schwann cell lines, pNF cell lines and MPNST cell lines. According to the results, we recommend using two reference genes (PPIA and TBP) in combination for gene expression analyses in MPNST related researches and using single reference genes (PPIA or PRLP0) alone or in combination in pNF related studies.

## Acknowledgments

Grateful acknowledgment is made to Prof. Jilong Yang and Prof. Vincent Keng, who kindly granted MPNST cell lines.

